# Thermal plasticity of the circadian clock is under nuclear and cytoplasmic control in wild barley

**DOI:** 10.1101/330829

**Authors:** Eyal Bdolach, Manas Ranjan Prusty, Adi Faigenboim-Doron, Tanya Filichkin, Laura Helgerson, Karl J Schmid, Stephan Greiner, Eyal Fridman

**Author notes:** Author for correspondence: Eyal Fridman Institute of Plant Sciences, Agricultural Research Organization (ARO), The Volcani Center, P. O. Box 6, 5025001, Bet Dagan, IsraelE-mail.

## Abstract

Temperature compensation, expressed as the ability to maintain clock characteristics (mainly period) in face of temperature changes, is considered a key feature of circadian clock systems. In this study, we explore the genetic basis for circadian clock plasticity under high temperatures by utilizing a new doubled haploid (DH) population derived from two reciprocal *Hordeum vulgare* sps. *spontaneum* hybrids genotypes (crosses between B1K-50-04 and B1K-09-07). Genotyping by sequencing of DH lines indicated a rich recombination landscape, with minor fixation (less than 8%), for one of the parental alleles, yet with prevalent and varied segregation distortion across seven barley chromosomes. Phenotyping was conducted with a high-throughput platform under optimal and high temperature environments. Genetic analysis, which included QxE and binary-threshold models, identified a significant influence of the maternal organelle genome (the plasmotype), as well as several nuclear quantitative trait loci (QTL), on clock phenotypes (free-running period and amplitude). Moreover, it showed the differential contribution of cytoplasmic genome clock rhythm buffering against high temperature. Resequencing of the parental chloroplast indicated the presence of several candidate genes underlying these significant effects. This first reported plasmotype-driven clock plasticity paves the way for identifying an hitherto unknown impact of nuclear and plasmotype variations on clock robustness and on plant adaptation to changing environments.

**Highlight:** Circadian clock robustness to high temperature is controlled by nuclear and plasmotype quantitative trait loci in a wild barley *(Hordeum vulgare* ssp. *spontaneum)* reciprocal doubled haploid population.

## Introduction

Radiation and adaptation of plant populations bear consequences on the genetic make-up of both nuclear and organellar genomes, driven by selective forces as well as stochastic genetic drift and founder effects. One of the great challenges in evolutionary biology is to distinguish between neutral and causal genetic variations that participate in the relationship between genetic and phenotypic variation. Moreover, we still need to realize the role of phenotypic plasticity in maintaining genetic divergence between populations. Identification of the genetic network regulating phenotypic plasticity will be key to deciphering mechanisms underlying, and following the trajectory of the respective alleles across habitat ranges will aid to understanding the adaptation history of a species (Price et al., 2003).

Circadian clock rhythm, which controls the pace and diurnal timing of plant development, physiology and biochemistry, is considered a central adaptive trait in sessile plants (Dodd et al., 2005)(Greenham and McClung, 2015). Temperature compensation, i.e. ability to maintain clock characteristics (period and amplitude) in face of temperature changes, is considered a key feature of circadian clock systems (Johnson et al., 2003). Indeed, analysis of molecular mutant phenotypes demonstrated that CCA and LHY function in buffering the clock under low and high temperatures, respectively (Gould et al., 2006). However, Arabidopsis plants speed their clocks in response to high temperatures (Edwards et al., 2005,Kusakina et al., 2014), with accession-specific variations noted in the degree of plasticity. For example, although *CCA1* and *LHY* transcription period shorten considerably across all Arabidopsis accessions in the transition from 17°C to 27°C, temperature effects on *LHY* expression period were shown varied between accessions (Kusakina et al., 2014). In addition, analysis of Arabidopsis responses to elevated temperature under controlled experimental conditions, indicated the presence of naturally occurring inter-accession variation in the buffering capacity against high temperature (Kusakina et al., 2014), which may represent adaptation to different temperature regimes in the original niche of the species. Yet, attempts to find an association between genetic variation of CCA and LHY, as well as seven additional clock genes, and temperature-dependent modulation of growth (as a proxy for fitness), failed (Kusakina et al., 2014). These results indicate genes outside the core clock machinery govern this relationship between buffering of the clock and DNA variation (Edwards et al., 2005) (Kusakina et al., 2014). One possible overlooked source for natural variation underlying buffering of the clock against temperature changes is the cytoplasmic genomes (plastid and mitochondria).

A recent work demonstrated that variation in the organellar genome can contribute to phenotypic or metabolomic differences between individuals, although the relative contribution of organellar versus nuclear genes remains unclear (Joseph et al., 2013). Effects of the cytoplasm on plant fitness and agronomic traits were first described by the German botanist Carl Correns (1909), and later in many plant systems (Roux et al., 2016)(Greiner and Bock, 2013). For example, disease resistance can be influenced by the genotype of the cytoplasm, i.e., the plasmotype, as can be yield and grain quality (Frei et al., 2003)(Sanetomo and Gebhardt, 2015). These traits are likely the result of local adaption of the original wild alleles, as shown for bread wheat *(Trictium aestivum*), where cytoplasmic influence on fruit quality is influenced by genotype-by-environment interactions (Ekiz et al., 1998)(Frei et al., 2003). Vis-à-vis regulation of the circadian clock rhythm, it was shown that mutations in nuclear-encoded chloroplast proteins (GUN1), known to be involved in retrograde signaling, or in mRNA maturation in the chloroplast, feed back into the nucleus and affect the expression of both circadian clock oscillator (e.g. CCA1) and output genes (CAB) (Hassidim et al., 2007). It is thought that photosynthetic electron transport generates a retrograde signal that adjusts the alternative splicing of nuclear-encoded transcripts encoding an SR protein (a regulator of RNA splicing) and other splicing factors (Petrillo et al., 2014). The nature of the signal and whether it is circadian-regulated is unknown, but circadian regulation of photosynthesis within chloroplasts has been suggested to cause circadian modulation of a retrograde signal that alters nuclear mRNA splicing (Dodd et al., 2015).

As part of our attempt to understand the factors driving genetic and phenotypic variations in the B1K population (Hubner et al., 2009)(Hübner et al., 2012; Hübner et al., 2013) (Bedada et al., 2014), including the significance of organelle genome variation, we generated a wild barley doubled haploid (DH) population. The 121 homozygous barley lines originated from two reciprocal F1 hybrids between two B1K accessions with distinct phenology and phenotypic plasticity. Here, we compared between the DNA diversity and phenotypic plasticity of the circadian clock rhythm among the DH population. This was done by genotype-by-sequencing and by examination of circadian clock free running period and amplitude under two temperature regimes (optimal and high) using the F method (Dakhiya et al., 2017) in a new high-throughput phenomics platform (SensyPAM). Quantitative genomic analysis with several genetic models allowed us to examine both the contribution of nuclear and organellar DNA variations to clock rhythm robustness against high temperature.

## Materials and Methods

### Plant material

One hundred and twenty one DH lines were derived from the F1 generation of the cross between B1K-50-04 and B1K-09-07, both of which are single-seed descents from the original Barley1K collection (Hubner et al., 2009). Spikes taken from two derived F2 plants were used to generate DH, using the anther culture protocol described by Cistué et al. (Cistué et al., 2003). Eighty-one DHs were derived from an F1 in which the female donor was B1K-50-04, and the remaining 40 were derived from the reciprocal hybrid (Table S1). The 121 DHs, referred to as the Ashkelon-Hermon (ASHER) population, are available to the research community upon request.

### Read mapping, SNP discovery, genotype calling and filtering for nuclear loci

Raw fastq files were processed with the standalone TASSEL 5 GBS v2 pipeline (Glaubitz et al., 2014) preparation for alignment and SNP discovery. Prior to the alignment steps, the reads were trimmed to 64 bases and then collapsed into a set of unique sequence tags. Short reads were mapped on the *Hordeum vulgare* genome assembly reference v2, available in the Gramene database (ftp://ftp.gramene.org/pub/gramene/release-56/fasta/hordeum_vulgare/dna/) using BWA 0.7.12-r1039 (Li and Durbin, 2009), with default parameters. SNP discovery and production were performed with the standalone TASSEL-GBS. Only uniquely aligned reads were considered for SNP calling. Genotype calling was carried out with the TASSEL-GBS plugins ‘DiscoverySNPCallerPluginV2’ and ‘ProductionSNPCallerPluginV2’, with minimum minor allele frequencyof 0.01.Perl scripts used to filter the 60,845SNP markers resulting from TASSEL-GBS pipeline (Table S2). Due to the homozygous nature of DH populations, we discarded all markers with heterozygotes parents from the analysis and only included those with parents with different genotypes. We also excluded markers exhibiting more than 2 alleles in the whole dataset (resulted with 2,327 SNPs; Table S3; Fig. 1).

**Fig. 1.**
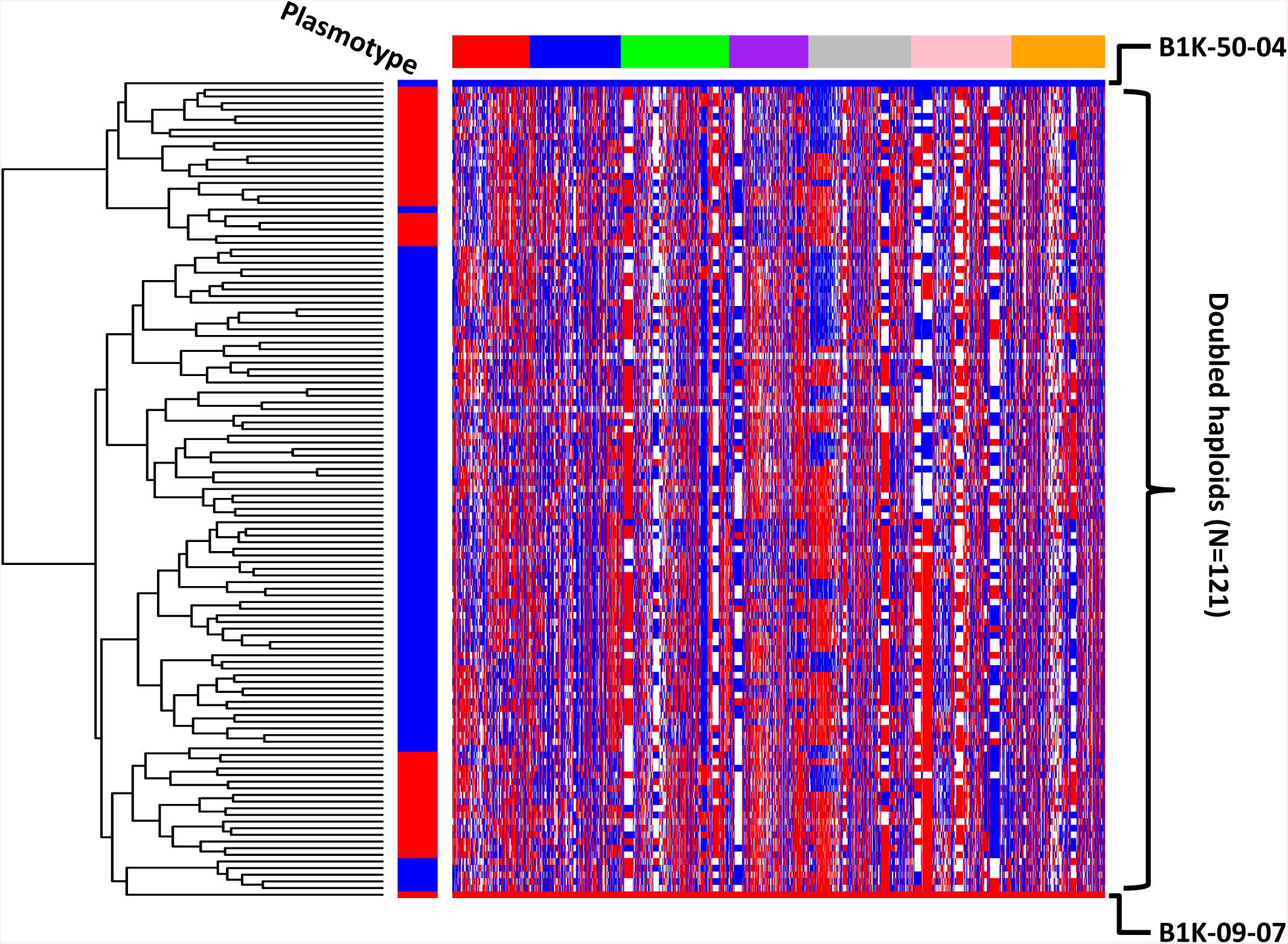
Graphical presentation of allelic distribution of 2237 markers (columns) and 121 doubled haploids (DH) lines (rows). The DH descended from two single reciprocal F_2_ genotypes derived from a cross between B1K-50-04 (blue) and B1K-09-07 (red). Plasmotype identity is color-coded.

For QTL mapping, four filtering criteria were applied to select a core set of markers: 1) heterozygous SNPs were omitted, since this is not expected in a DH population, 2) The SNP was scored in ≥90 DHs, 3) a minimal allele frequency of 10% among the DHs, 4) adjunct SNPs with identical pattern across DH were represented by only one of them. This procedure yielded a list of 760 high-confidence nuclear SNPs used to characterize the genetic make-up and allow QTL mapping (Table S3). Furthermore, prior to QTL mapping, a linkage map was calculated to position the 760 markers on the 7 barley chromosomes (Fig. S1). In addition, to determine possible allelic distortion, a Chi-square test was performed against the null hypothesis of a 1:1 ratio between the A and B alleles in each locus.

### Plastid genome (plastome) sequencing and SNP discovery

Illumina paired-end libraries (375 bp insert size) of total barley DNA were used to sequence the plastid genomes of the parental lines, B1K-50-04 and B1K-09-07. For this, 250-bp reads were mapped against the barley reference plastomes of the cultivar Barke (KC912687; (Middleton et al., 2014)), using the SeqMan NGen v.14.1.0 (DNASTAR, Madison, WI, USA) software. Larger insertions/deletions were identified by paired-end mapping. The plastid DNA contigs (comprising of LSC, IR_B_ and SSC, with an average coverage of 234 and 331, respectively) were annotated using GeSeq v.1.20 (Tillich et al., 2017) and SNPs called in SeqMan Pro v15.0.1 (DNASTAR, Madison,WI, USA).

### Circadian clock phenotype of doubled haploids under optimal and high temperature

To mimic the natural growth conditions of the wild barley population in the Southern Levant, where the original Barley1K infrastructure was collected (Hubner et al., 2009), we modified an earlier protocol used for free running clock measurements (Dakhiya et al., 2017). Mainly, instead of growing the seedling under a regime of 16 h light and 8 h dark, plants were grown to the emergence of the fourth leaf under 8h light and 16h dark, at a constant temperature of 20°C. Following this entrainment of the plants for app. four weeks, they were moved to a high-throughput SensyPAM (SensyTIV, Aviel, Israel) custom-designed to allow F measurements in up to 240 plants for each experiment. The SensyPAM includes a large carousel suitable for 14 plastic trays (Fig. S2A) that spins the trays one at a time between different compartments including continuous light, dark adaptation, imaging chamber and post-imaging compartments (Fig. S2C). Each tray is filled with water and can hold up to 24 plants that are placed along the sides of the tray and the fully expanded upper leaves are pinned down to a black sponge. The imaging chamber is equipped with an acA-1920-40um CCD camera (Basler, Exton, PA, US), with a 647nm long-pass edge filter (Semrock, Rochester, New York, US) and a LED light panel. The LEDs are royal blue (450 nm), red-orange (630 nm), far-red (740 nm) and cool white (450-700 nm) (REBEL type, Lumiled, San Jose, CA, US). The trays are circling every five min. on a carousel between continuous light, dark adaptation, imaging and post imaging compartments. Before the tray is photographed, it is kept in a dark adaptation chamber for two steps (10 min.) (Fig. S2B), after which, it is kept for three more min. at the imaging chamber under dark (total 13 min. dark adaptation; Fig. S2C), followed by the PAM timeline (Table S5). To identify the individual plants, 3-4 areas of interest (AOI) for each leaf are marked in the SensyPAM software (the number of AOI depends on the size of the leaf) before the experiment begins (Fig. S2D). The light pulses and durations are similar to those applied in the saturation pulse method (Schreiber, 2004), with saturating flashes of 1600 μimoll m^2^ s^-1^ and actinic light of 457 μimoll m^2^ s^-1^ (example output in Fig. S2E). The F parameters, and light durations are explained in Table S5. The SensyPAM is located in a room with controlled temperature and light. In the ASHER DH population, plants were grown under optimal temperature (OT, 22°C) or high temperature (HT, 32°C) for several cycles of experiments (Fig. S3), For the clock measurement, F was measured every 2.5 hours, for 3 days, in light. Each genotype was grown twice under each temperature, with 4 plants included in each round. For the clock analysis, mean NPQlss ((Fm-Fmlss)/Fmlss) was calculated and the period, amplitude and relative amplitude error (RAE) were extracted using the BioDare2 website (https://biodare2.ed.ac.uk) (Fig. S2F). Input Data “cubic dtr” and Analysis Method “MFourFit” (Zielinski et al., 2014). To assess the robustness of these rhythms at high temperatures, the delta of the FRP and AMP for the two temperatures (HT-OT) were compared. In a different approach, the FRP means between the two treatments was compared for each DH line using the t Student’s test to infer a binary variable, i.e. if the difference was significant (p<0.01), the response was 1, otherwise, it was 0. This approach takes into account the variance in each line per temperature.

### Genome-wide and cytonuclear interaction QTL analysis

To further dissect the genomic architecture underlying circadian clock robustness against high temperature in the DH population, a QTL analysis was conducted using three different approaches. Initially, we performed a single QTL analysis with the subtraction (delta) between HT and OT for clock FRP and AMP values (dFRP and dAMP, respectively). This also included a di-genic analysis assessing the possible interaction between nuclear and cytoplasmic loci. In addition, to identify loci associated with trait plasticity, a genetic model containing the QxE component was formulated and probability (P value) for this interaction with the temperature (QxE) was calculated for specific loci (Sasaki et al., 2015). Last, we hypothesized that the signal triggering a change in the clock rhythm, may work under a *threshold* model (Falconer, 1965). Under such a model, an unobservable quantitative variable, termed *liability*, underlies the binary phenotype, with a significant change in the clock rhythm elicited at above-threshold concentrations, and insignificant or no changes at below-threshold liability values (Falconer 1965). We therefore translated the FRP phenotype of the DH lines into a binary vector (see above). This allowed performance of a genome scan for loci that may switch the system from heat compensated [maintaining the same rhythm; (Johnson et al., 2003)] to a responsive one, i.e. one with a positive or negative clock rhythm change.

The genome-wide and di-genic (nuclear and cytonuclear) QTL interaction analysis of the DH population for different traits was carried out using inclusive composite interval mapping (ICIM; (Li et al., 2007)) with the IciMapping V4.1(Meng et al., 2015) software package. IciM 4.1 uses an improved algorithm of composite interval mapping for biparental population. It has a high detection power, reduced false detection rate and less biased estimates of the QTL effects, which minimize the bias for small population sizes. Initially, a BIN function was used to remove the redundant markers. A linkage map was constructed using the MAP function, with the steps (parameters) as grouping (by anchor order), ordering (by input), and rippling (with SARF, window size = 5) and using Kosambi mapping function (Fig. S1). The output was then used to find the QTLs for the different traits, using the BIP function. For the additive analysis, and digenic epistasis interaction, mapping with the parameters was performed: missing phenotype = deletion, step = 1 cM, and probability in stepwise regression = 0.001. To identify significant QTLs, a LOD score of 2.5 was manually set at a significance level of p < 0.05. A more stringent analysis obtained the LOD score by performing a 1000-permutation test. The digenic epistatis interactions were estimated at both LOD values of (2.5) and (5). The QTL by environment interaction (QxE) was also assessed with the inclusive composite interval mapping (ICIM) method, using the MET function of the software QTL IciMapping 4.1 (Li et al., 2007, Meng et al., 2015). To study the cytonuclear interaction, the cytoplasm genome was considered as an extra chromosome (plasmotype), in addition to the 7 nuclear chromosomes of barley, and was used for the QTL and epitasis analysis.

## Results

### Nuclear and plastid genome variation within the ASHER wild barley doubled haploid population

To explore a potential link between genetic diversity and circadian clock robustness to changing environments, 121 homozygous DHs were generated from a reciprocal cross between B1K-09-07 and B1K-50-04 (Table S1; see Methods). The two accessions originated from two different habitats of wild barley in the Southern Levant, the most allele-rich part of the *H. spontaneum* gene pool (Pankin et al., 2018). Illumina genotype, determined by sequencing the 121 lines and comparing them to the DNA of the parental lines, yielded c. 52 million reads, out of which, on average, 81.8 % matched the *Hordeum vulgare* genome assembly reference v2 (ftp://ftp.gramene.org/pub/gramene/release-56/fasta/hordeum_vulgare/dna/)(Table S2). Phylogenetic analysis of the DHs showed clearly that the DH descended from each F1 plant clustered together (Fig. 1). The genome-wide allele frequency was then analyzed to examine the possibility that selection (in F1 or F2 generations), or genetic drift (in F2 generation) occurred during process (Cloutier et al., 1995). Figure 2A and Table S3 depict the segregation ratios across seven barley chromosomes among the 121 DH lines, for a total of 2,237 markers that were different between two parental lines. In total, 7.6% or 7.4% of the markers drifted to allele frequencies higher than 0.9 for the B1K-50-04 or B1K-09-07 allele, respectively (hereafter coded as ‘A’ or ‘B’ alleles for any locus) suggesting differential gamete selection (Nixon, 2006). This pattern implies that the genomic landscape could cover a significant portion of the genome (app. 85%) for identifying causal variation for phenotype of interest since mean of each genotypic group will be calculated from at least 12 DHs. Nevertheless, transmission ratio distortion (TRD), or significant bias from the expected 1:1 ratio between the two parental alleles in the DH progeny, was prevalent in both directions. The rate of significant TRD (Chi-test, p<8*10E-5, corrected to multiple testing) ranged from 10% of markers in chromosome 7 to 90% for chromosome 4 (Fig. 2B; Table S3).

**Fig. 2.**
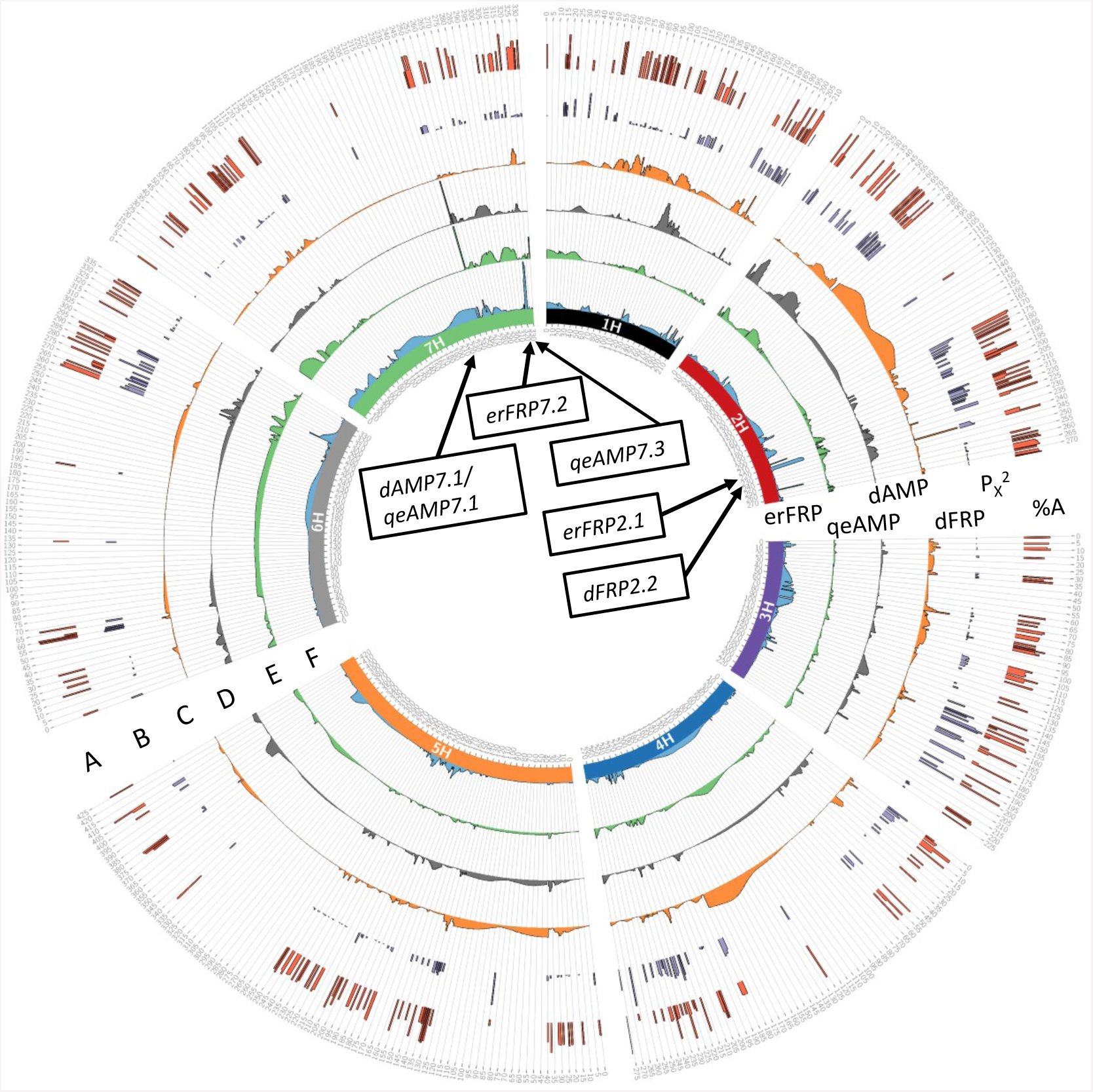
Circos plot of the allele frequency and QTL analysis results for the DH population. (A) Allele segregation ratios depicted with percentage of the A allele (%A), and (B) statistically tested with Chi-square tests (-logioP) for distortion from expected 1:1 ratio for the 760 loci. QTL results for (C) dFRP (delta free-running period), (D) dAMP (delta amplitude) (E) qeFRP (QTL by environment interaction) and (F) erFRP (environmental response of FRP), binary-threshold model (see Methods). Barley chromosomes in the Circos plot are depicted in different colors in the inner circle. For each trait, the track represents the QTL LOD score in a 1-cM window. Window positions (in cM) are ordered clockwise, per chromosome. The maximum heights of the y-axis are 89.6 [%], 17.0 [-LogP], 3.13 [LOD], 3.1 [LOD], 5.27[LOD], and 3.86[LOD] for graphs A to F, respectively. QTLs are marked with arrows and the names are indicated in the inner circle (see Table S6 for details).

### Carriers of the B1K-50-04plasmotype show increased robustness of the circadian rhythm

To explore the potential link between cytoplasm identity (chloroplast and mitochondria) and circadian rhythm, 114 DHs were examined in a high-throughput manner for circadian clock rhythm by scaling-up the chlorophyll F method (Dakhiya et al., 2017) (see Methods; Fig. S2). In this analysis we asked whether plastid or mitochondria organelles (together referred to as plasmotype) could be involved in regulating the pace of the clock, as well as robustness of its characteristics.

Overall, FRP values ranged varied between 21.5-31.7 h under OT, but showed a significantly narrower range under HT (21.0-25.9 h (Fig. 3A)). The mean FRP of the DH population under the two temperatures differed significantly (Student’s t-Test; p<0.0001), i.e. clock rhythm accelerated from 25.3 ±2.3 h under OT to 23.2±1.02h under HT (Fig. 3B I). The difference between the FRP of the two sub-populations was highly significant under OT (P<0.0001), but not under HT (P=0.11; Fig. 3B II). More specifically, under OT, the mean FRP of population 0950 was 26.5 ±2.2 h, and circadian clock was significantly accelerated (shortened FRP) to 23.03 ±1.0 h under HT (Fig. 3B III). In contrast, mean FRP of subpopulation 5009 was 24.7 ±2.05 h in OT and was slightly accelerated (23.35 ±1.0 h) under HT (Fig. 3B III). In summary, there was a difference of more than 2 h (2.2) in the delta of the clock FRP (dFRP) between the carriers of the different cytoplasms. A significant difference in the dFRP, i.e. the difference between mean FRP under HT versus OT, between the carriers of the two plasmotypes was noted (Student’s t-test; p<0.0001). Taken together, these results implicate that the maternal genome of 5009 DHs confers greater circadian clock robustness or less plasticity to HT.

**Fig. 3.**
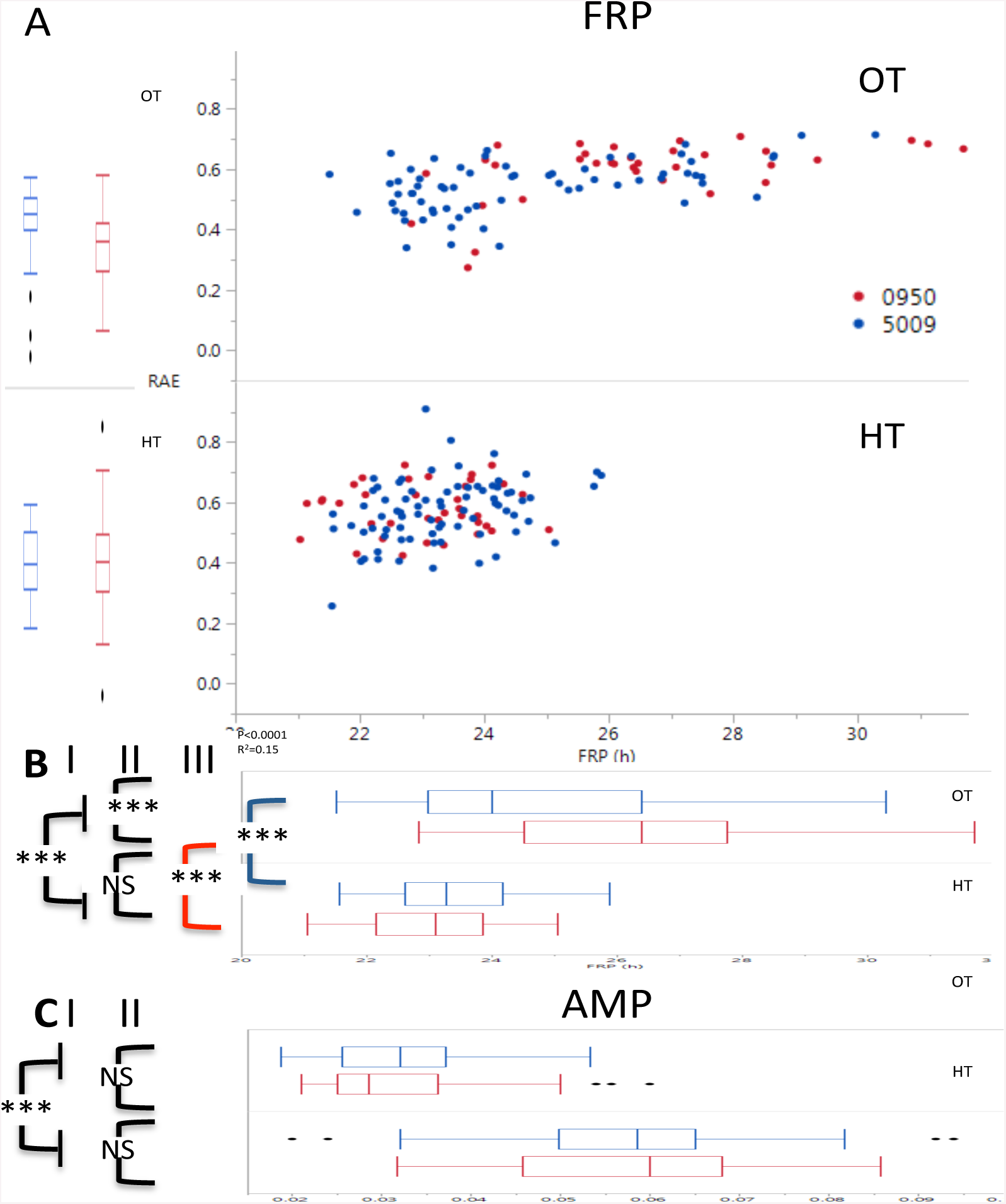
Circadian clock rhythm in DH plants is differentially shortened under high temperature between carriers of the B1K-09-08 compared to B1K-50-04 plasmotype. (A) Free running period (FRP) vs. relative amplitude error (RAE) of 114 DH lines originating from two F_2_ reciprocal wild barley, i.e. carrying either B1K-09-07 (0950; red) and or B1K-50-04 (5009; blue) plasmotype, under two treatments; optimal temperature (OT) at 22°C and high temperature (HT) at 32°C. Mean difference between treatments for the entire population (I), between plasmotypes 0950 and 5009 under OT and HT (II) and between OT and HT in each subpopulation (0950 and 5009) (III) were determined for (B) FRP and (C) amplitude (AMP). For each T-test, the p value is depicted as ***: P<0.001 or NS: not significant.

Clock amplitude (AMP) showed a general and significant (P<0.001) doubling between high and lower temperature (0.032 in OT and 0.058 in HT; Fig. 3C I). However, under both temperatures there was no significant difference between two cytoplasm sub-populations (Fig. 3C II), implicating a neutral effect of the cytoplasmic source on the amplitude.

### Single-locus analysis of nuclear variation and circadian clock robustness

*Single QTL analysis of the delta between HT and OT:* Three QTLs were identified for dFRP, when setting a manual input of LOD =2.5, which is a relatively permissive threshold (data not shown). However, when increasing the stringency of the genetic analysis using 1000 permutations (LOD = 3.29), a single significant and major QTL was identified on chromosome 2 (flanking markers S2H_ 718,338,890 and S2H_722,401,229). Segregation in this locus explains the 13.4% variation recorded for the trait, and with additive effect of+1.0 h (the positive value corresponds to allele A vs. B and represents half of the difference between two homozygous genotypes; Fig. 2C; Table S6). The relatively low number of significant QTLs underlying the clock plasticity between HT and OT environments, was also evident for the dAMP; only a single locus on chromosome 7 (flanking markers S2H_498,472,330and S2H_ 510,903,725), which explained 11.15% of variation for the trait, was identified (Fig. 2D; Table S6).We next considered the possible contribution of di-genic epistasis interactions to the overall variation of FRP and AMP. Di-genic QTL analysis (see Methods) at low stringency (LOD=2.5) identified fifteen pairs of interacting loci, including one in tight vicinity to the single QTL on chromosome 2 (flanked by S2H_706,464,739 and S2H_713,705,096; Table S7). Increasing the stringency to LOD=5, left only two significant di-genic interactions, albeit with PVE values below 4% (Table S7). Setting a similar stringent threshold of LOD5 for the digenic interactions underlying variation in dAMP, failed to yield a significant pair of interacting loci. Those di-genic interactions found with a LOD2.5 explained each less than 3.5% of the total phenotypic variation (Table S7). Notably, no significant cytonuclear interactions between the cytoplasm and the nuclear loci were identified.

*Markers by environment interaction (QxE) analysis of clock traits:* Since the choice of method used to map trait plasticity has significant consequences on the results (Josephs, 2018), we used the same dataset of DH clock phenotypes to specifically map loci underlying QxE. The analysis showed only a partial overlap with genome scans of the delta for both traits between environments. For the clock AMP, the major QTL on chromosome 7 (flanking markers S7H_ 498,472,330 and S7H_ 510,903,725) was also identified, at a higher LOD score (5.27), and its segregation was explaining large portion of the phenotypic variation (PVE=36.43%). In addition, the QxE QTL analysis identified another locus on same chromosome, between S7H_ 651,765,311 and S7H_656,04,8080 (Fig 2E; Table S6). In contrast, this QxE analysis failed to identify neither significant QTL on any chromosome location, nor any association between FRP variation and cytoplasm source.

*A threshold-binary model analysis:* Our different approach for the QTL analysis stemmed from the hypothesis of a *threshold* mechanism (see Methods) and therefore the phenotype of each DH was pre-considered and converted to a categorical binary vector. For each DH line, comparison of FRP means between the two groups of plants grown under two treatments (t Student’s test, p<0.01) showed that only 12 out of 66 DH lines (18.5%) were significantly different for 5009 samples, including a negative delta for some lines, while 15 out of 34 (44.1%) 0950 DHs showed accelerated clock FRP under heat (Fig. S4; Fig. 4A). These significant differences (Pearson P>ChiSq=0.004) between two sub-populations are in agreement with the overall robustness of the 5009 vs 0950 DH. This “single locus” analysis of the plasmotype effect was then extended to a genome-wide scan of loci affecting the non-random distribution of the binary trait between genotypic groups of each marker (A or B; see Methods). This analysis also yielded two QTLs on chromosomes 2 and 7, but at locations different from loci identified for QxE or the delta of the traits (Fig. 2F; Table S6). The one QTL on chromosome 2, flanked between S2H_698,875,542 and S2H_702,308,910, showed a significant (Pearson P>ChiSq=0.0004) non-random distribution of the binary phenotype between two genotypic groups, with carriers of the ‘A’ (B1K-50-09) included 81.6% non-affected, as compared to half of that (40%) among carriers of the ‘B’ (B1K-09-07) allele (Fig. 4B). Similarly, only 14.8% of carriers of the ‘A’ allele at the chromosome 7 QTL changed their FRP under HT compared to OT (Fig. 4C). This division between affected and non-affected DH lines was significantly different than the 38.7% affected among B-carriers at this locus (Pearson p>ChiSq=0.014).

**Fig. 4.**
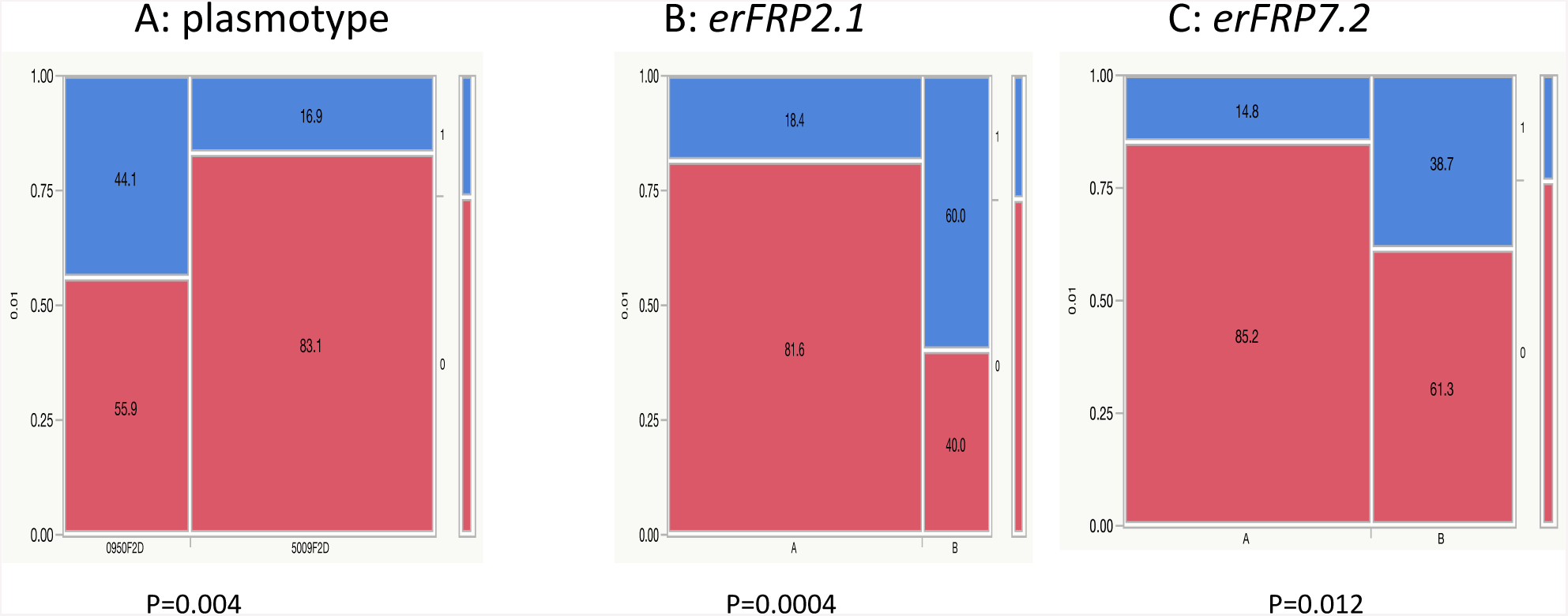
Distribution of the binary environment-responsive FRP trait (erFRP) between plasmotype and QTL genotypes. (A) Proportion of the affected DH population (scored as ‘1’), i.e., those that show a significant difference (student’s t-test; P<0.01) in FRP under HT vs OT, was significantly higher (**χ^2^** test) in carriers of the B1K-09-07 compared to B1K-50-04 cytoplasm. Similarly, clock rhythm of DH carrying the B1K-50-04 allele in (B) *erFRP2.1* or (C) *erFRP7.2* loci was significantly less affected (scored as ‘0’) as those with the B1K-09-07 cytoplasm. Pearson P value of **χ^2^** test for each locus is indicated below.

## Discussion

### Automated PAM as a phenomics tool

Given plants’ high levels of morphological plasticity, compared to animals, crop phenomics presents distinct challenges (Tardieu et al., 2017). Here, we present a new phenomic tool that allows for non-invasive measurement of the circadian clock rhythm under optimal and high temperatures for a large set of genotypes in a relatively short time and with high repeatability. This is crucial for QTL mapping and for development of models able to disentangle and simulate plant behavior in a range of habitats. Until very recently, circadian clock rhythm was measured by analysis of expression of genes belonging to the core clock complex, e.g., CCA and LHY (Gould et al., 2006). This created an obstacle since plants had to be harvested at each time point and therefore, either wounded plants were repeatedly sampled (possibly changing the natural condition of the measured plant) or required growth of each genotype in large numbers to allow statistical sensitivity The F method, which requires dark adaptation, poses some limitations for small labs due to relatively high costs of engineering and manufacturing of the LED panel and CCD camera. Possible alternatives include either replacement or supplementation of the CCD camera with a thermal camera that can capture the leaf temperature. Canopy temperature is known to follow aperture of the stomata, which are also under the control of the circadian clock (Kreps and Kay, 1997). However, one possible caveat of this measurment is the differences and possible independence between the circadian rhythm in stomata cells and the rest of the leaf (Yakir et al., 2011). Further studies are therefore required to make such comparison between F and thermal based measurements and perhaps combine the two to provide either a “uniform”, organ- or cell-specific rhythms.

### Contribution ofplasmotype to circadian clock rhythm and plasticity

In the analysis of the clock characteristics between the two reciprocal sub-populations, we noted a significant difference in the rhythm and its robustness under changing temperature regimes (Fig. 3). These findings indicate a role of either the chloroplast or mitochondria in clock regulation, and demonstrate the utility of the DH population for dissection of the interplay between nuclei and organelles with regard to clock regulation. Most of our knowledge of such interplay relates to anterograde (nucleus-to-plastid) control of the organelle, and less to retrograde (plastid-to-nucleus) signaling. The latter is thought to be controlled by the metabolic state of the chloroplast, including sugar status (Rolland and Sheen, 2005)(Haydon et al., 2013) and reactive oxygen species, which are, like sugars, affected in part by photosynthesis, in general, and high light (Vandenabeele et al., 2004) or by high temperature, in particular. Interestingly, mutation in an RNA binding protein destined to the chloroplast, resulted in altered circadian rhythms (Hassidim et al., 2007). Nevertheless, the nature of the signal and sensitivity to circadian rhythm are unknown. Of note, it has been suggested that circadian regulation of photosynthesis within chloroplasts causes modulation of a retrograde signal that alters nuclear mRNA (Dodd et al., 2015), including alternative splicing of nuclear-encoded transcripts SR protein (a regulator of RNA splicing) and other splicing factors (Petrillo et al., 2014). However, there are no reports of circadian clock plasticity sensors or initiators within the chloroplast.

Sequencing of the plastids (see Methods) identified 81 SNPs between the plastid genomes of B1K-50-04 and B1K-09-07, including 40 singletons and 42 SNP indels, ranging in size between one and eight nucleotides (Table S4). Notably, the number of polymorphisms in coding regions between the parental lines (six non-synonymous and seven synonymous variants), exceeded the variation identified in earlier studies (Middleton et al., 2014). However, more careful analyses of sequence variation in the barley plastome should include harmonization of gene annotations. The six non-synonymous mutations found between the organelle genomes of two donors suggest a relatively low number of candidate genes, if coding regions are considered the sole factor linking genome source to circadian clock rhythm. Future studies will be required to test if one of these underlies the significant effects of the cytoplasm on circadian clock rhythm. Notably one of the identified non-synonymous mutations resided within the rpoC1 gene, which is part of the PEP protein complex. It is composed of subunits encoded by both plastid (rpoA, rpoB, rpoC1, and rpoC2) and nuclear genes (sig1-6) and coordinated rates of molecular evolution between plastome and nuclear encoded genes were observed, at least in Geraniaceae (Zhang et al., 2015). PEP itself has a broad effect on the plastid transcriptome. PEP-controlled genes drive rhythms of transcription from the blue light-responsive promoter of chloroplast-encoded *psbD* (Noordally et al., 2013) and potentially of other chloroplast genes. Therefore, communicating timing information to the chloroplast genome is proposed to act as a circadian signal (Noordally et al., 2013)(Dodd et al., 2015). Nevertheless, validating such causal variations *inplanta* is conditioned on advancing our ability to achieve plastid transformation in higher plants, which is nevertheless challenging for monocots (Yu et al., 2017). In addition, despite the relatively low known sequence variation between barley mitochondrial genomes (only 3 SNP, between *H. vulgare* and *H. spontaneum*, including 2 in intergenic regions, out of a genome of 525,599 bp) (Hisano et al., 2016), we are yet to compare the sequence variation between parental lines before determining whether variation in this organelle genome underlies differential clock phenotypes among DH sub-populations.

### Genetic and biochemical basis for circadian clock plasticity

Flowering time in barley and in other model plants is an example of a research field that gained extensive knowledge from integration of qualitative and quantitative genetics, which assisted in building models to explain the genetics and biochemistry underlying variation between plant populations (see at wikipathways.org/index.php?query=flowering&title=Special%3ASearchPathways&d oSearch=1&sa=Search). When taking the QTL approach, rather than forward genetics of mutants, was taken, the trait is treated as a quantitative variable and association is made between allelic and trait variations, which is usually was validated by additional mutants in candidate genes (Comadran et al., 2012). Taking the classic QTL approach in our study led to identification of very few QTLs that were associated with plasticity or response of the circadian rhythm to high temperature (Fig. 2). This may be the result of presence of only two possible alleles in our system, as opposed to multiple alleles found in recently studied multi-parent populations (Maurer et al., 2015) (Merchuk-Ovnat et al., 2018). Nevertheless, the QTL found for FRP and AMP plasticity do not seem to fall on candidate genes in the barley gene pool, which were proposed as potential regulators of this trait under domestication (Pankin et al., 2018). Thus, we may have identified, beyond the novel connection to organelle-dependent causal variation, some hitherto unknown loci that regulate clock plasticity under high temperature.

Furthermore, we propose to adopt threshold models, which were traditionally considered for binary disease traits with a polygenic basis (Falconer, 1965)(Xu and Atchley, 1996), to dissect circadian clock plasticity as a binary trait. The main difference from disease phenotypes is the population angle, i.e. in the present case, the binary score is given based on statistical difference between several individuals of each genotypic group (marker/plasmotype). The biological hypothesis behind this approach is that circadian clock-modulating signaling in response to elevated temperature, acts as a semi-conductor, with amplification of only signals above a certain threshold, then translated to a change in the circadian rhythm. From a statistical point of view, it adds more confidence since the binary score is based on several individuals. Implying this binary approach has identified sets of loci distinct from those identified with conventional QTL mapping; in these loci, including the plasmotype, a significant non-random distribution of the affected and non-affected genotypes was found between the two genotypic groups.

## CONCLUSIONS

This high-throughput circadian clock analysis of a newly developed reciprocal barley doubled population, indicated a hitherto unknown relationship between plasmotype variation and thermal phenotypic robustness. Additional studies with nearly isogenic lines for the “regular” and “binary” QTL, will enable examination of the stability of these loci across temperature gradients and in different genetic backgrounds, with the long-term goal of identifying the biochemical elements underlying allelic differences and their causality. Ongoing work is aiming to examine the consequences and relationships between clock phenotypes and whole-plant fitness phenotypes under common garden conditions. This integrated analysis will question the importance of circadian clock rhythm plasticity versus stability in plant fitness under fluctuating environments.

## Acknowledgements

We wish to thank Royi Levav, Oded Anner and Elad Lifshin (SensyTIV, Israel) for the valuable design, construction and implementation of the SensyPAM hardware and software. The technical assistance of Khaled Bishara and Inbar Anner is highly appreciated. The discussions and technical assistance of Prof. Rachel Green and Yuri Dakhiya (The Hebrew University) were instrumental in setting up the high-content phenotyping system. We acknowledge the editing of the manuscript by Yehudith Posen. The FACCE-ERA-NET (Grant # 406/14) and ISF 1270/17 grants to E.F supported this research. Work by SG was supported by the Max Planck Society.

## Contributions

E.B and E.F. designed the experiments, collected, analyzed and interpreted data, and wrote the manuscript. E.B., A.F. and M.R.P. were involved in the data analyses, their interpretation and in writing the manuscript. T.F. and L.H. generated the doubled haploid population from crosses made by E.F. K.A.S. extracted SNPs from GBS data with assistance from A.F. S.G. sequenced and analyzed the chloroplast genomes.

